# Translation variation across genetic backgrounds reveals a post-transcriptional buffering signature in yeast

**DOI:** 10.1101/2023.11.28.568778

**Authors:** Elie M. Teyssonniere, Yuichi Shichino, Anne Friedrich, Shintaro Iwasaki, Joseph Schacherer

**Author notes:** Corresponding authors (S.I.) and (J.S.). first co-authors (Elie M. Teyssonniere, Yuichi Shichino).

## Abstract

Gene expression is known to vary among individuals, and this variability can impact the phenotypic diversity observed in natural populations. While the transcriptome and proteome have been extensively studied, little is known about the translation process itself. Here, we therefore performed ribosome and transcriptomic profiling on a genetically and ecologically diverse set of natural isolates of the *Saccharomyces cerevisiae* yeast. Interestingly, we found that the Euclidean distances between each profile and the expression fold changes in each pairwise isolate comparison were higher at the transcriptomic level. This observation clearly indicates that the transcriptional variation observed in the different isolates is buffered through a phenomenon known as post-transcriptional buffering at the translation level. Furthermore, this phenomenon seemed to have a specific signature by preferentially affecting essential genes as well as genes involved in complex-forming proteins, and low transcribed genes. We also explored the translation of the *S. cerevisiae* pangenome and found that the accessory genes related to introgression events displayed similar transcription and translation levels as the core genome. By contrast, genes acquired through horizontal gene transfer events tended to be less efficiently translated. Together, our results highlight both the extent and signature of the post-transcriptional buffering.

## Introduction

Transcript and protein abundance variations are well-known sources of phenotypic diversity across individuals. Protein abundance is influenced by both transcriptional and post-transcriptional regulations, which ultimately affect the final phenotypes. There are several cellular mechanisms involved in the modulation of final protein abundance, including mRNA stability, translation initiation and protein degradation (1). In the last decades, various technologies have greatly facilitated the detailed exploration of all these steps. Some of these technologies include DNA high-throughput sequencing methods such as RNA sequencing (2), and mass spectrometry (3), which enable a global description of transcriptomic and proteomic dynamics. By associating these data with genomic data, we can greatly improve our understanding of the mRNA and protein abundance regulation at the population level (4–8). Furthermore, the core of the translational process can be precisely dissected with the development of ribosome profiling (or Ribo-Seq) (9, 10). This strategy relies on the sequencing of mRNA fragments covered by the ribosomes during the translation process, revealing which parts of the transcriptome are actively being translated. This method primarily allows quantification of translational activity for each mRNA (number of reads corresponding to one mRNA), but can also be used to examine ribosome density along the mRNA, revealing translational characteristics such as ribosome stalling or elongation rate (11).

The budding yeast *Saccharomyces cerevisiae* has been a powerful model for ribosome profiling experiments, as this technique was developed on this organism (10). Translational variation in yeast has been explored with ribosome profiling on several occasions (12–17). Interestingly, several of these studies highlighted that the transcriptional variations tended to be buffered when looking at the translational variations (13–16). This phenomenon is known as post-transcriptional buffering and has also been observed when comparing transcriptomic and proteomic datasets (18, 19). However, despite recurring observations, the mechanisms underlying this post-transcriptional buffering are still poorly understood. Moreover, while transcription and protein abundance have been extensively monitored, translation itself has been considerably less studied, and no clear description of the phenomenon has been made at this level. More globally, translational variation remains largely unexplored, and several known sources of expression variation, such as accessory ORFs (Open Reading Frames), have yet to be investigated at the translational layer.

Here, we conducted ribosome profiling and RNA sequencing in the same conditions on eight *S. cerevisiae* natural isolates coming from very diverse ecological environments and being genetically different (20). We first compared the transcriptional and translational variations, and found that they had similar functional patterns. Metabolism-related genes tended to be more variable across the eight isolates while essential genes and genes involved in molecular complexes had more conserved transcription and translation regulation. Interestingly, we found that the transcriptional profiles were less correlated to each other compared to the translational profiles. Accordingly, Euclidean distances and expression variations (quantified using the absolute log2 transformed foldchanges for each gene in each isolate pairwise comparisons) were significantly higher in the transcriptomic data. We also found that, overall, almost half of the genes displaying transcriptional variation between isolates were affected by the post-transcriptional buffering, indicating that this phenomenon is a strong determinant of the translational variations. More importantly, we found that this phenomenon has a specific signature in terms of affected genes. We observed that essential genes and protein complex-related genes as well as lowly transcribed genes tended to be preferentially buffered. Furthermore, we investigated the transcription and translation of accessory Open Reading Frames (ORFs) present in the eight isolates, particularly those acquired through introgression or horizontal gene transfer (HGT) events. We observed that introgression-related ORFs were similarly transcribed and translated compared to their orthologs, while HGT-related ORFs displayed a significantly lower translation efficiency than the rest of genes. Together, our results provide an overview of translational variations as well as an accurate description of post-transcriptional buffering.

## Results

### Ribosome profiling and RNA sequencing across eight natural isolates

We performed both RNA sequencing (RNA-seq) and ribosome profiling (Ribo-seq) in replicates on eight genetically diverse *S. cerevisiae* isolates (Table S1), which were cultivated and harvested in the exact same condition (See Material and Methods). These isolates were selected to represent the genetic diversity of the species (Figure S1) and were grown on a synthetic complete medium. The genomes of all isolates were all previously sequenced (20), and in addition to their very different genetic backgrounds, they also came from very diverse environments.

For RNA-seq and Ribo-seq analyses, raw counts were normalized using the anota2seq algorithm (TMM-log2 approach) (21), allowing to study a total of 3, 200 genes (see Methods). Correlations between the RNA-seq and Ribo-seq replicates are consistently high (Figure S2A-B). Moreover, replicates of the RNA-seq and Ribo-seq profiles of each isolate well cluster using PCA analysis (Figure S3A-B). Interestingly, we observed a strong correlation between RNA-seq and Ribo-seq data from each sample (Spearman correlation test between 0.52 and 0.80), highlighting the relationship between the level of transcription and translation (Figure S4). To gain an overall view of intraspecific variation, we performed strain-pairwise Spearman correlation tests on the two datasets (Figure 1A). RNA-seq and Ribo-seq correlation matrices showed similar patterns, indicating once again that transcriptional variations were largely reflected at the translational level. Consistently, we observed a strong positive correlation between the coefficients of the two correlation matrices (Figure 1B).

**Figure 1.**
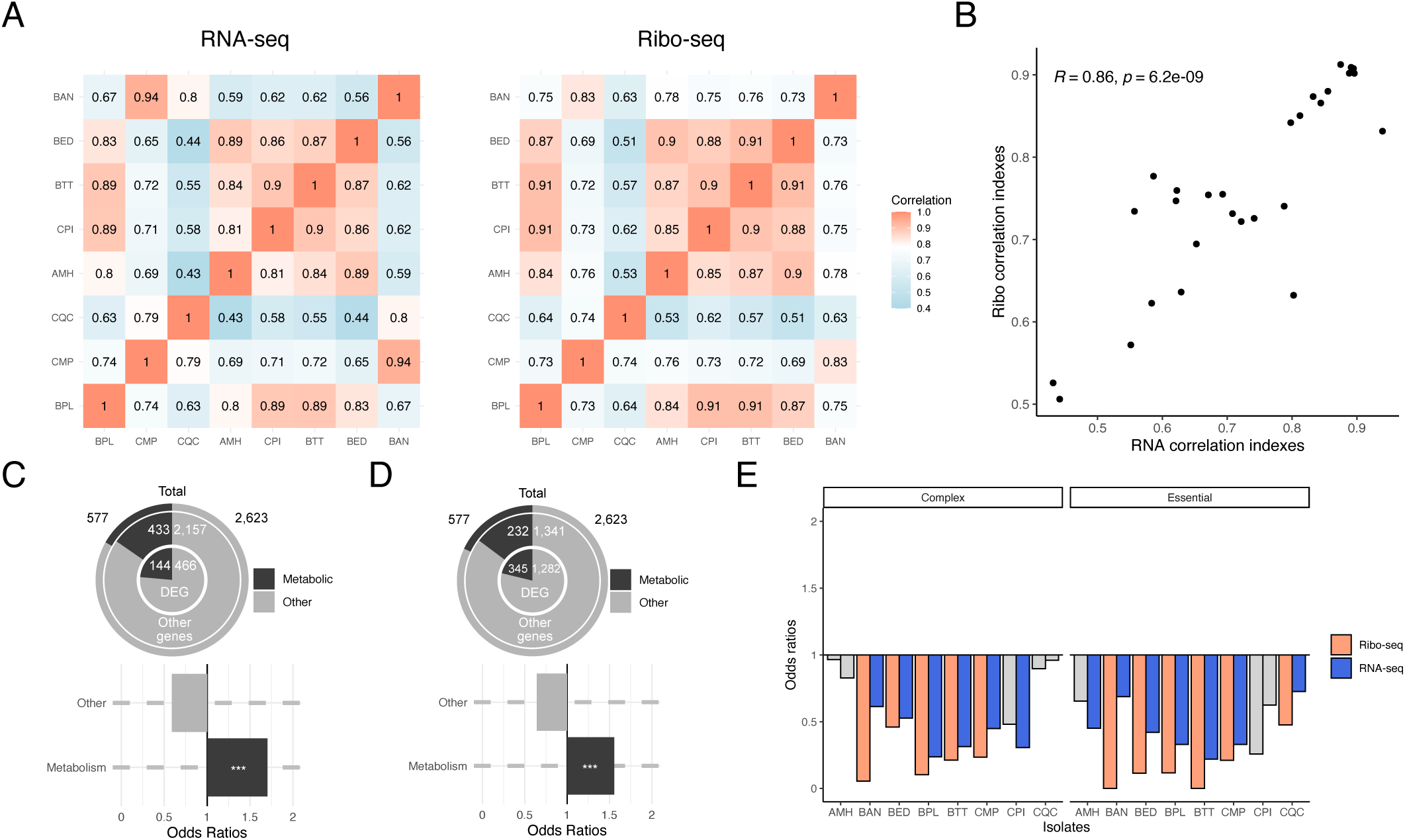
Exploration of the transcriptional and translational variations. (A) Correlation matrices (Spearman correlation test) of each RNA-seq and Ribo-seq isolate pairwise comparison (all the coefficients displayed significant P-values). (B) Correlation between the RNA-seq and Ribo-seq correlation matrices. (C, D) Enrichment of metabolism-related genes in the (C) Ribo-seq and (D) RNA-seq DEG. Both enrichments were significant (FET, respective P-value: 1×10^-4^ and 2.1×10^-6^). (E) Enrichment of protein complex related genes and essential genes among the set of DEGs for each isolate. The grey bar corresponds to unsignificant enrichments.

Next, we sought to identify genes that did not follow these correlation trends in order to detect genes with variable regulation at both levels (transcription and translation). We therefore performed a differentially expressed gene (DEG) detection using the *limma* R package (22, 23). Briefly, we compared the expression level of each gene for each isolate at both levels relative to the other 7 isolates (see Methods). We identified a total of 3, 429 DEGs (with 1, 627 unique genes) and 816 DEGs (with 610 unique genes) in the RNA-seq and Ribo-seq datasets, respectively (Table S2). The number of DEGs varied greatly depending on the isolates, as it ranged 143 and 1, 136 for the RNA-seq (median = 328) and from 17 to 533 for Ribo-seq (median = 34) (Figure S5A-B). The CQC isolate shows the highest number of DEGs, consistent with that this strain having the most different transcriptome and translatome profiles (Figure 1A and Figure S3).

To investigate functional enrichment among these identified genes, we conducted a Gene Ontology analysis (GO) on the two sets of DEGs (24, 25). We found similar results for both expression levels with a significant enrichment of terms related to metabolism and respiration (Table S3). To confirm the overall enrichment of metabolism genes, we concatenated all GO terms associated with metabolism (see Methods) to generate an overall “Metabolism” gene group and looked at whether DEGs were enriched among it. We still detected a significant enrichment for RNA-seq and Ribo-seq DEGs (Figure 1C-D). This observation can be explained by the fact that the eight isolates used in this study were obtained in distinct environments (Table S1) and could have adapted their regulation of several metabolic functions to different trophic conditions. It also emphasizes that transcriptional and translational variations from one individual to another are functionally related.

In contrast, we found that both sets of DEGs (transcription and translation) were overall depleted of essential and protein complex related genes (26–28) (Figure 1E). Interestingly, depletion was less pronounced in the Ribo-Seq data, suggesting that these genes were less likely to exhibit variable regulation at the translational level compared to the transcriptional level (Figure S6).

Together, our results highlight that expression variation is unequal among the genes. While metabolism related genes display important transcriptional and translational variations among the eight isolates, essential genes and protein complex related genes are related to a lower transcriptional and translational variation.

### Post-transcriptional buffering at the translation level across isolates

Despite similar trends and functional enrichments of DEGs, we observed lower variability at the translational level compared to the transcriptional level. By comparing the correlation matrices (Figure 1A), the Ribo-seq correlation coefficients were indeed significantly higher than those of RNA-seq (Figure 2A). We could confirm this lower variability by examining the Euclidean distance between isolates using either normalized RNA-seq or Ribo-seq data (Figure 2B). Distances between strains were significantly higher at the transcriptional level, suggesting more similar translational profiles. We also quantified the variation in gene expression at both levels by computing the absolute value of the log_2_ transformed foldchange (*|log2(FC)|*) for each gene in a pairwise comparison (n = 89, 600) (see Methods). We found that *|log2(FC)|* was significantly and approximately 10% higher in RNA-seq data (mean = 0.22) compared to Ribo-seq data (mean = 0.20) (Figure S7), suggesting that the transcriptional variation tends to be overall buffered.

**Figure 2.**
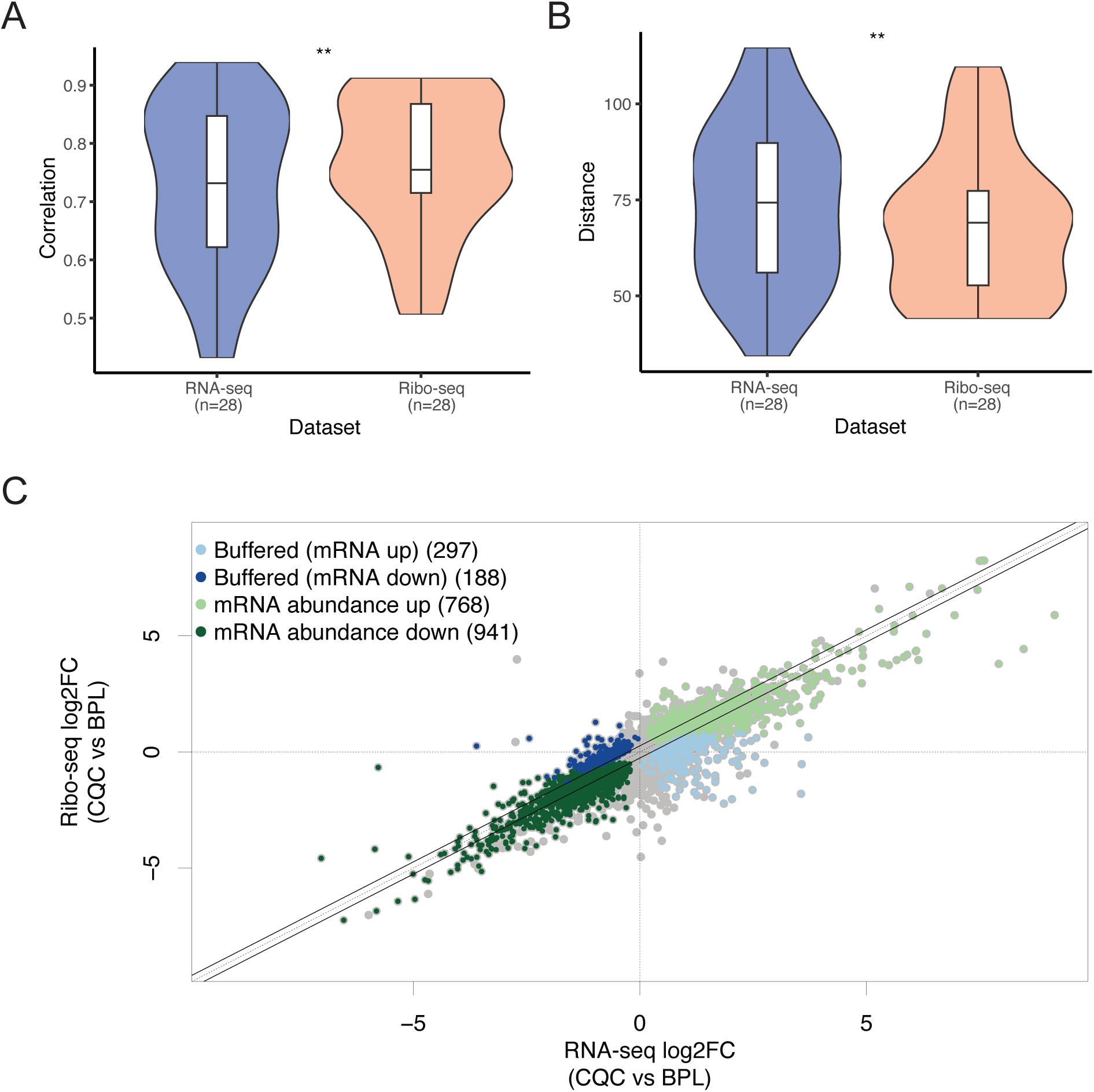
Post-transcriptional buffering at the translation level. (A) Isolate pairwise correlation levels (from figure 1A) differences between RNA-seq or ribo-seq data. The correlations coefficients were significantly higher in Ribo-seq data (paired Wilcoxon test P-value = 0.0022). (B) Differences between the RNA-seq and Ribo-seq Euclidean. The distances are significantly higher in RNA-seq data (paired Wilcoxon test P-value = 0.0041). (C) Example of comparison between RNA-seq log2(FC) and Ribo-seq log2(FC) as generated by Anota2seq in the pairwise comparison of CQC vs. BPL isolate. The blue dots correspond to the genes where an increase (in light blue) or decrease (in dark blue) in mRNA levels in the CQC vs. BPL isolates is buffered at the translational level. Conversely, the light and dark green dots correspond to the genes where the increase or decrease in mRNA levels is conserved at the translational level.

All these results are consistent with the phenomenon of post-transcriptional buffering that has been observed previously (13–16). We sought to determine the extent to which this phenomenon affected the transcriptional variation in each pairwise comparison. We therefore used the *anota2seq* R package (21) in order to analyze the transcriptional and translational variation in each pairwise comparison. In fact, this analysis enabled us to determine which genes exhibit transcriptional variation, and which genes are buffered at the translational level (Figure 2C). The number of buffered genes reached an average of 768 genes per pairwise comparison, ranging from 349 to 1, 222 genes (Table S4, Figure S8A). Overall, this set of buffered genes represents an average of 44.8% of the genes with transcriptional variation, ranging from 21.6% and 81.5% (Figure S8B).

Overall, almost half of the genes showing transcriptional variation between two isolates are affected by post-transcriptional buffering, suggesting that this phenomenon is widespread and strongly influences translational variation.

Taken together, our results suggest that although transcriptional and translational variation are similar in pattern and function, their strength is strongly attenuated at the translation level because of the post-transcriptional buffering phenomenon.

### Signature of the post-transcriptional buffering at the translation level

Despite several observations of post-transcriptional buffering, this phenomenon remains largely unknown, especially at the functional level (13–16). We sought to further characterize the general rules underlying this phenomenon by looking for genes that would be preferentially affected by the post-transcriptional buffering at the translational level.

With that in mind, we used the set of genes previously detected as buffered in each pairwise comparison (Figure 2C, Table S4) and looked for a functional signature. We therefore used GSEA (29) using the *fgsea* R package (see Methods) (30) and found that buffered genes were more likely to be associated to certain central functions for the cell, such as cell cycle, transcription regulation, chromosome segregation or mRNA transport (Table S5).

Seeing this enrichment of central cell function in the buffered genes, we then focused on the content of essential (27, 28) and protein complex-related genes (26). We checked in each pairwise comparison whether the buffered genes were enriched in these two groups of genes. We found that the buffered genes were enriched in protein-complex related genes compared to genes with unbuffered transcriptional variation (Figure 3A). Similarly, essential genes were also enriched among the buffered genes (Figure 3B). Consistently, we found that the odds ratios obtained from the Fisher’s exact tests (FET) used to calculate the enrichment in either essential or complex-related genes were significantly higher than 1 (Figure S9). Together, these results support the fact that the genes are unequally affected by the post-transcriptional buffering at the translational level, with essential genes and protein complex-related genes preferentially buffered.

**Figure 3.**
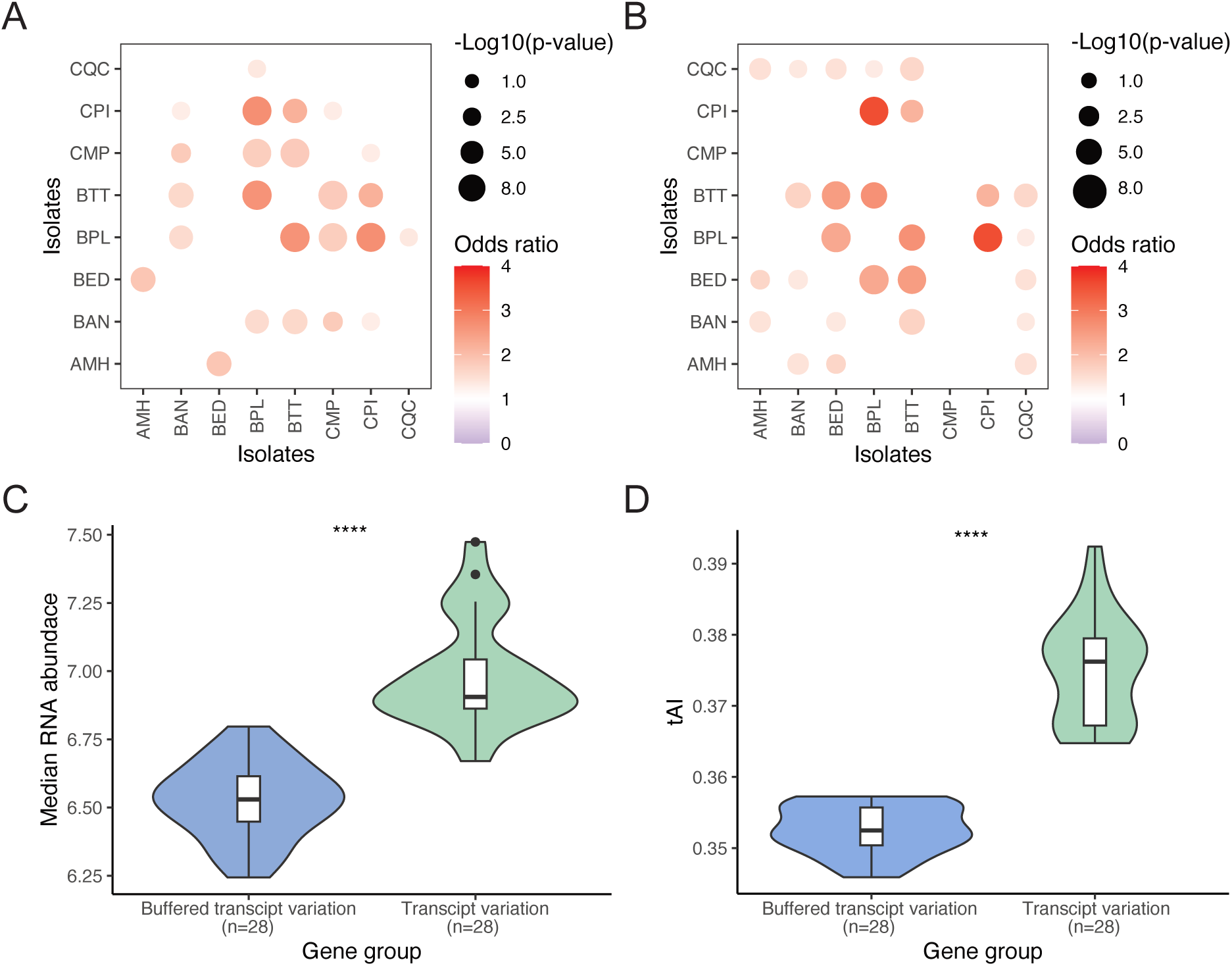
Post-transcriptional buffering has a specific signature. (A, B) Enrichment test in each of the pairwise comparisons for (A) protein complex-related genes and (B) essential genes. The size of the dot corresponds to the -log10(P-value) obtained using Fisher’s exact tests and the color corresponds to the estimated odds ratios. Pairwise comparisons without any dot correspond to the cases where no significant enrichment or depletion could be detected. (C, D) Average (D) RNA-seq and (C) tAI levels of the buffered genes versus the genes with unbuffered transcription variation (respective Wilcoxon test P-value: 1.6×10^-14^ and 1×10^-15^).

Interestingly, essentiality and protein-protein interactions are features that have been debated for their influence on protein sequence evolution (31–34). Another major determinant of sequence evolution is the gene expression level, as highly expressed proteins are known to be more conserved (34–37). We therefore sought to determine whether the transcriptional level was also involved in preferential buffering. Surprisingly, we found that the buffered genes were significantly less transcribed (average mRNA level = 6.53) than genes with unbuffered transcript variation (average mRNA level = 6.91) (Figure 3C). Consistently, when examining each pairwise comparison individually, we observed that in most of the cases (24 out of 28), buffered genes are less transcribed than the genes with unbuffered transcript variation (Figure S10). We also observed that low mRNA levels of buffered genes remained true when compared to all other genes (Figure S11). We suspected that this difference might be an artifact of the fact that low-expressed genes have a biased biological signal-to-noise ratio, and therefore the transcriptome variation is artificially increased for low abundant mRNA. However, we found no correlation between mRNA level and variance (Figure S12), supporting the reliability of the mRNA difference between buffered and unbuffered genes. Taken together, these results highlight that transcription level is also a determinant of the post-transcriptional buffering phenomenon, since buffered genes tend to be less transcribed.

We sought to confirm these results by exploring the codon usage bias of the buffered genes since this feature is known to be related to expression level (38, 39) and essentiality (40). We computed a codon usage bias index for each gene using tRNA Adaptation Index (tAI) (41, 42), and we confirmed that this index overall correlated with RNA-seq or Ribo-seq data (Table S6). We then compared the average tAI values of the buffered group to those of the unbuffered group. We observed a significantly lower tAI in the buffered group (median tAI = 0.35) compared to the unbuffered group (median tAI = 0.38) (Figure 3D). Similar results were observed when performing the comparison in each isolate pairwise comparison (Figure S13). This observation supported the previous results of expression level difference between the two groups.

Overall, these results highlight the fact that the phenomenon of post-transcriptional buffering preferentially affects essential, protein complex-related genes, or genes with lower transcription levels and therefore has a specific signature. This behavior toward some specific categories of genes has never been shown before and highlights evolutionary constraints affecting the translational regulation of these genes.

### Transcription and translation variation of accessory genes

Recent advances in *S. cerevisiae* population genomics have highlighted the presence of more than 1, 700 variable ORFs (accessory genes) in this species (20). Our translation exploration across multiple individuals from very different genetic and environmental origins is a unique opportunity to explore the translation of such ORFs. In our eight isolates, the number of these ORFs varies between 63 to 215, corresponding to a total of 446 unique accessory ORFs (median = 94 accessory ORFs per strain), but depending on the isolate, between 53% and 89% of them were expressed (Figure S14). Our eight isolates exhibit variable profiles in terms of accessory ORFs origins (Figure S14). Two strains differed notably to the others in their compositions: the CPI isolate due to a very high number of ORFs acquired by introgression and the BPL isolate due to genes acquired through horizontal gene transfer (HGT).

Regarding the CPI isolate, this strain was originally isolated in Mexico and has been described as part of the “Mexican agave clade” (20). This subpopulation has a high number of introgressed ORFs coming from the yeast *Saccharomyces paradoxus* (median = 161 ORFs per strain vs 25.75 in the overall population). The CPI isolate has 131 expressed ORFs coming from introgression events, and 124 ORFs had known orthologs in S288C. In order to strictly explore the impact on transcription and translation, we focused on 13 out of the 124 ORFs that were homozygous for the *S. paradoxus* allele in the CPI isolate and found no expression difference between these ORFs and their orthologs in the 7 other strains (again at the transcriptional and translational level) (Figure 4A-B). These results imply that the transcriptional and translational regulation of ORFs acquired by introgressions from *S. paradoxus* is similar to their regulation of their orthologs.

**Figure 4.**
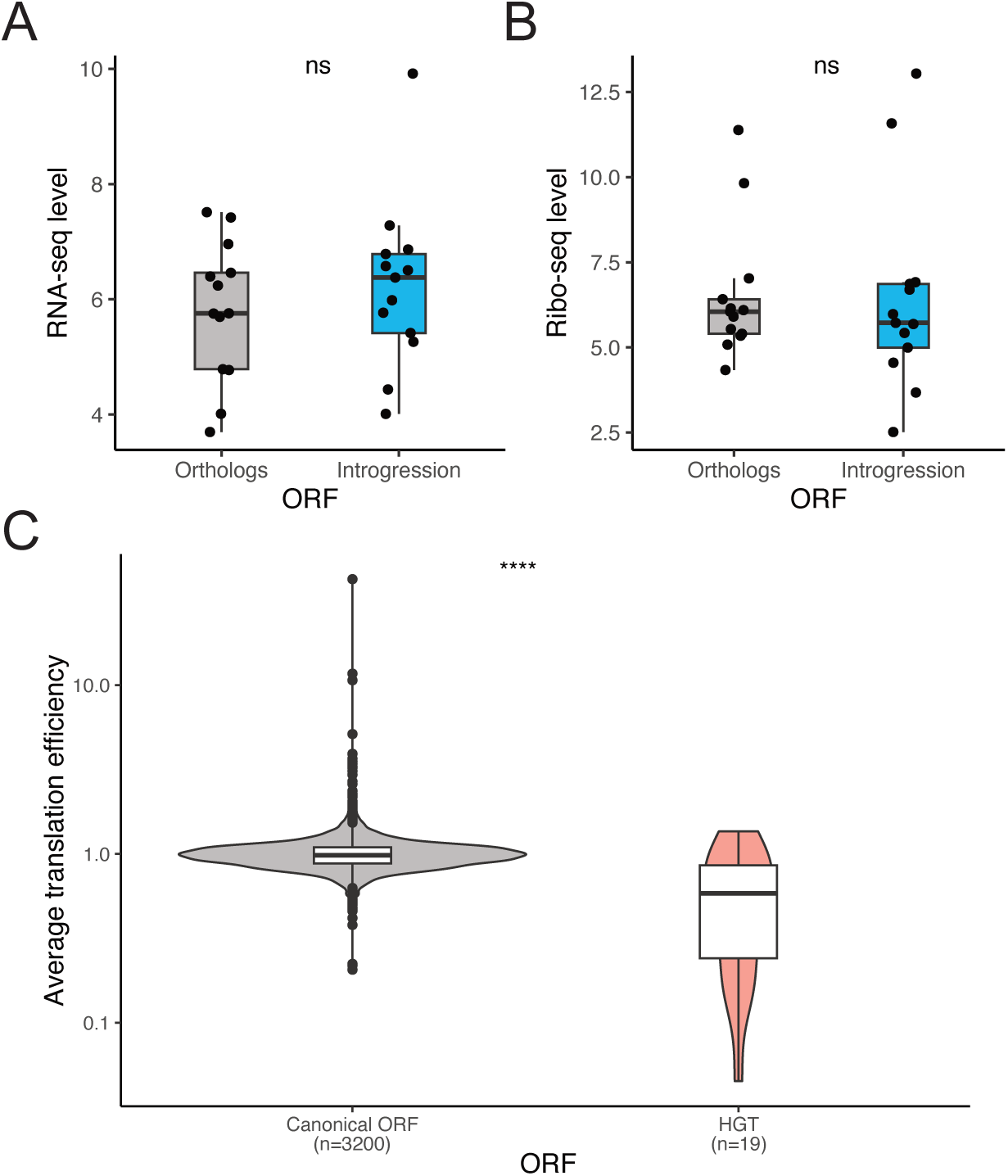
Translation levels of the *S. cerevisiae* pangenome. (A, B) RNA-seq and Ribo-seq level of the ORFs acquired through introgression event in the CPI isolate and being homozygous for the *S. paradoxus* allele (n = 13) and their orthologs in the other isolates. No difference in term of transcription of translation were observed between the introgressed ORFs and their orthologs. (C) TE difference between the ORFs acquired through HGT in the BPL isolate and the other ORFs (Wilcoxon test P-value = 0.0002).

We then focused on the expression of the 19 accessory ORFs coming from HGT in the case of the BPL isolate (Table S7). This strain has already been described as part of a wine subpopulation (20) and the occurrence of HGT events in this type of strain has already been observed (43, 44). Briefly, the coexistence of *S. cerevisiae* with other yeast species in the wine environment led to gene transfer that can confer evolutionary advantage in the winemaking environment. We compare the expression of these ORFs to other genes in the BPL isolate (Figure S15A). Surprisingly, while the overall mRNA levels of the HGT ORFs were similar to the rest of the ORFs, their translation did not follow this trend and was much lower than the other ORFs. We then compared the translation efficiency (*i.e.*, the ratio between the translation level and the transcription level of a gene) of the HGT ORFs with the rest of the genes and found a significant difference where HGT ORFs were less efficiently translated than other genes in the BPL isolate (mean HGT TE = 0.58, mean HGT other genes = 0.95) (Figure 4C). A possible explanation for this would be that HGT might be less adapted to the translation machinery or regulation of *S. cerevisiae* since these ORFs are from genetically distant yeast species (43, 44). We tested whether these ORFs differ in terms of codon usage bias using tAI as described above. We found no significant differences (Figure S15B), suggesting that the difference in translation origins is due to something else. Taken together, our results suggest that the translational behavior of accessory genes is highly dependent on their origin.

## Discussion

Translational variation is a major determinant of the transcriptome-proteome relationship, and therefore plays a central role in the phenotypic diversity observed in natural populations. However, the translation process itself remains largely unexplored, and the central mechanisms driving translation variations are still poorly understood. In this study, we have precisely monitored translation variations across natural isolates of *S. cerevisiae* using ribosome profiling.

Gene expression is known to differ across individual. We observed that the translational and transcriptional variations were similar in terms of trends and functions, with metabolism-related genes displaying the greatest variation while the translation regulation of essential genes and protein complex-related genes were more conserved across individuals. These results are consistent with recent large-scale exploration of mRNA abundance (45) and highlight that gene expression plasticity might be driven by the metabolism preferences between isolates coming from very different environments (46). Conversly, expression variation of genes with central and essential functions is likely to be deleterious and tend to be therefore more conserved (47).

Our dataset also allowed to get better insight into the phenomenon of post-transcriptional buffering at the translational level (13–16). As suggested by several analyses of our dataset (*e.g.*, Euclidean distances, isolate pairwise correlation level, |log2(FC)|), we found lower variability in gene expression at the translational level. Taken together, these results clearly indicate the presence of post-transcriptional buffering in our dataset. Overall, we found that, almost half of the genes (44.8%) on average showing transcriptional variation are affected by this phenomenon, highlighting the central role it plays in shaping translational variation across individuals.

Interestingly, we found that post-transcriptional buffering has a specific signature. It preferentially affects genes that are essential or related to protein complexes, as well as genes with low transcript levels. Several reasons could underlie this preferential buffering. For complex-forming protein, it is well established that the imbalance of complex components can be deleterious for (48–50), partly for stoichiometric reasons (51). More generally, protein complex genes are known to have stronger regulatory control at the protein level rather than at the mRNA level (52) and programmed translation of complex components precisely proportional to stoichiometry was not only found in yeast (53), but also in bacteria and plants (54–57). Essential genes are known to carry a central and highly conserved function in the cell (26), which can lead to higher constraints on gene expression evolution (58). However, the conditional nature of essentiality (59, 60) and the ongoing debate on the importance of essentiality on evolutionary constraint (31, 32) suggest that the link between expression conservation and essentiality remains unclear. Regarding the fact that buffered genes tend to be less transcribed than other genes, this is surprising since very abundant proteins are usually well conserved (34–37). Thus, we would have expected that the regulation of highly expressed genes would also be highly conserved as well, and therefore preferentially affected by the phenomenon of post-transcriptional buffering.

The cellular mechanisms underlying post-transcriptional buffering are still poorly understood. At the proteomic level, several hypotheses have been proposed to partially explain the phenomenon, such as autoregulation or degradation of excess protein complex members (1, 19, 61–63). However, these findings are limited to specific types of genes and are therefore unlikely to explain the widespread nature of the phenomenon. Moreover, our results focus on buffering at the translational level, where hypotheses on the mechanistic basis of the phenomenon are even scarcer. Our results show that post-transcriptional buffering preferentially affects genes with very different characteristics (*e.g.*, essential genes and low-expressed genes), which could suggest that the mechanisms underlying the phenomenon are multiple. Translation initiation and regulation are obvious candidates, especially in yeast, where a systematic survey of post-transcriptional regulation has identified hundreds of regulators (64).

Finally, working with genetically distinct natural isolates allowed to explore the translation of a part of the *S. cerevisiae* accessory genome, which is something that has been barely investigated so far. Accessory genes had very different translation dynamics depending on their origins of acquisition. While introgressed ORFs displayed similar levels of translation compared to their orthologs, HGT-related ORF were less translated, resulting in low translation efficiency. Several reasons could explain this decrease in translation efficiency. As mentioned above, they could be less adapted to the translation regulation of *S. cerevisiae*, but we found no evidence for this. Another possibility is that these ORFs were acquired and maintained in winemaking environments. Our culture in complete medium is obviously a poor representation of such an ecological condition, therefore these ORFs might be less useful and thus their translation is downregulated. These results on the pangenome nevertheless remain limited due to the low representation of the entire pangenome of *S. cerevisiae* (20). More generally, a broader view of *S. cerevisiae* population translation would improve our understanding of the post-transcriptional buffering phenomenon.

Overall, our results highlight the importance of the post-transcriptional buffering at the translation level, as well as its specific signature. Moreover, they give one of the first insight into the translation dynamics of a specific part of the genome such as accessory genes.

## Materials and methods

### Strain, culture, and flash freezing

The complete list of isolates used in this study is available and described in Table S1. The strains were grown in liquid SC medium (Yeast Nitrogen Base with ammonium sulfate 6.7 g.l^−1^, MPbio, OH, USA; amino acid mixture 2 g.l^−1^, MPbio; glucose 20 g.l^−1^, Euromedex, France). The culture was maintained until the strains reached their growth mid log phase using an optical plate reader (Tecan infinite F200 pro). The cells were then filtered using 0.45 µm MCE membrane (Merk Millipore, France). The filters were then plunged into a 50 mL tube containing liquid nitrogen and stocked in a -80°C freezer before being used for ribosome profiling and RNA sequencing experiment.

### Ribosome profiling and RNA sequencing

The library preparation for ribosome profiling was performed as previously described with modifications (65, 66). Cells on the filters were mixed with frozen droplets of 600 µL lysis buffer (20 mM Tris-HCl pH 7.5, 150 mM NaCl, 5 mM MgCl_2_, 1 mM dithiothreitol, 100 µg/mL cycloheximide, and 1% Triton X-100) and crushed using Multi-beads Shocker (Yasui Kikai, Japan). Lysate containing 20 µg of total RNA was digested with 10 U of RNase I (Lucigen, WI, USA) for 45 min at 25°C. Ribosomes were precipitated by sucrose cushion and ultracentrifugation. RNAs ranging from 17 to 34 nt were excised from a polyacrylamide TBE-Urea gel. The rRNA was depleted using riboPOOLs Kit (*Saccharomyces cerevieiae*) (siTOOLs Biotech, Germany).

For RNA-seq, total RNA was purified using TRIzol LS reagent (Thermo Fisher Scientific) and Direct-zol RNA Miniprep Kit (Zymo research, CA, USA). Following the removal of rRNA by riboPOOLs Kit (*Saccharomyces cerevisiae*), the sequencing library was prepared with SEQuoia Express Stranded RNA Library Prep Kit (Bio-Rad, CA, USA). The ribosome profiling and RNA-Seq libraries were sequenced on a HiSeq X Ten platform (Illumina) with a pair-end 150 bp.

### Sequence data alignment, quantification, and normalization

Alignment and quantification of ribosome profiling and RNA-Seq data were performed as previously described with modifications (66). After the removal of the linker sequence and the splitting based on sample barcode, we removed reads that mapped to non-coding RNA (ncRNA) sequences using STAR 2.7.0a (67). Despite the fact that we used an rRNA depletion method for both RNA-seq and Ribo-seq, our libraries were highly contaminated with ncRNA, resulting in a relatively low reads number input for the alignment: between 127, 087 and 4, 464, 548 reads for Ribo-seq. Remaining reads were aligned to the S288C *S. cerevisiae* genome using STAR 2.7.0a (67). For the analysis of the accessory ORFs (open reading frames), the reads were also aligned to all the ORF detected in the pangenome of *S. cerevisiae* (20). The A-site offsets of ribosome footprints were determined according to the location of the 5′ end of reads mapped to start codons. For RNA-seq, offsets were set to 15 for all mRNA fragments. Reads corresponding to the first and last five codons of each coding sequence (CDS) were excluded from the analysis. The custom scripts will be available upon requests.

### Gene expression data normalization

We normalized the raw data (read count) using the standard built-in normalization of the *anota2seq* R package (21). Briefly, this consisted of a log2-TMM normalization. We used this normalized data in all subsequent analyses unless otherwise specified. In order to visualize the quality of our results, we performed both PCA and correlation matrix using the normalized replicate data. We also computed a mean expression value for each gene in each isolate that we used to construct the Spearman-based correlation matrix from Figure 1. Ultimately, after the exclusion of ORF without read in any samples, or ORF without expression variance, we ended with a list of 3, 200 genes for which both transcription and translation were quantified (Table S8, S9).

### Variable genes detection and analysis characteristics

We sought to detect genes that exhibited gene expression variation at both RNA-seq and ribo-seq levels. We did this by using the *limma* R package (22, 23) which compute a linear model for each genes in order to detect differentially expressed genes (DEG). We applied a one-vs-all strategy, where each isolate is compared to the 7 others, using their weighted mean expression. We considered a gene to be differentially expressed if the FDR-adjusted p-value as reported by *limma* was lower than 10^-5^.

We looked for functional enrichment in the DEG and performed gene ontology exploration (24, 25, 68) on the detected DEG using the *gprofiler2* R package (69). We also grouped all the GO terms associated to metabolism by selecting the terms labeled with the word “metabolic” and using this large gene group to detect any significant enrichment of metabolism related genes in the DEG using Fisher’s exact tests (FET). We also looked if we could find enrichment or depletion of essential gene or protein complex related genes. (26–28) in the DEG using Fisher’s exact tests (FET). Firstly, we questioned if the variable genes detected earlier displayed any enrichment or depletion of essential genes or genes part of protein complexes with FET.

### Detection of a post-transcriptional buffering phenomenon

In order to see if expression variation across the 8 isolates was stronger in RNA-seq or Ribo-seq data, we firstly computed the Euclidean distances between each strain using the normalized data from both datasets. We also compared the isolates pairwise correlation (displayed in figure 1) obtained with the Spearman correlation test. We finally quantified the expression variation (in both RNA-seq and Ribo-seq data) based on the log_2_(FC):

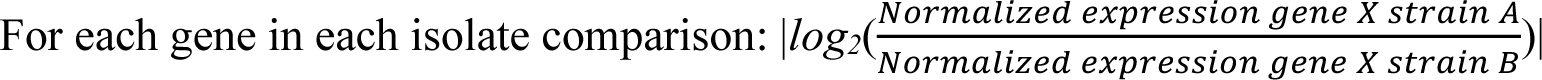

The more this value increases, the more variable is gene expression between to isolate for a specific gene.

We then sought to explore the extent of the phenomenon by looking at which genes were affected by post-transcriptional buffering. In other words, we wanted to determine for which transcriptional variation is buffered when looking at translational variation. We used the *anota2seq* R package (21) which compares the transcriptional and translational fold change in the comparison between two samples and allows to classify the genes in different groups (transcriptional variation, buffered variation, etc.) by analyzing the change in translational efficiency using analysis of partial variance. We used the *anota2seqRun* function on the normalized RNA-seq and ribo-seq data with the following parameter: onlyGroup = T. We performed this for each strain and for each of the 7 comparisons vs. the other isolates. For the later analyses, we focused only on the buffered genes and the gene with transcriptional variation as detected by *anota2seq*.

### Buffered and conserved regulation genes characteristics

We sought to further characterize post-transcriptional buffering by examining whether this phenomenon has a specific signature in terms of affected genes. We first looked at the number of times each gene was included in the buffered genes and used this measure to perform gene set enrichment analysis (GSEA) using the *fgsea* R package (30) and the GO terms as the annotation. We considered an annotation to be detected if its corrected P-value was less than 0.1. Annotations associated with preferential buffering were those with NES (new enrichment score) > 0.

Then, in each isolate pairwise comparison, we examined the enrichment of essential genes or protein complex-related genes among the buffered genes compared to the gene showing transcriptional variation between two isolates. We used FET for this. In addition, in each pairwise comparison, we calculated a mean mRNA level for the two sets of genes to compare the transcription level of these genes. We did the same using a measure of codon usage bias: tAI (see below for details on tAI calculation).

### Codon usage bias influence

We used the tAI (tRNA adaptation index) index (41, 42) to estimate the codon bias usage of each gene. Briefly, tAI is an index showing how much a gene is adapted to the tRNA genome structure in terms of codon usage. In this perspective, we first calculated the tRNA copy number of our 8 strains with tRNAscan-SE with default parameters (70, 71) using the assembled genome sequences from (20). Then, for each isolate, we used the tRNA copy number to compute the tAI using a Perl program available on https://github.com/mariodosreis/tai (41, 42) using default parameters. The resulting dataset was a tAI value for around 5, 600 genes (some genes were discarded during the calculation) in each isolate. Ultimately, we could calculate an overall tAI (mean of the 8 or less tAI values) for 3, 746 genes. For each isolate, we correlated the expression levels to the tAI values.

### Accessory ORF analysis

Using the mapping done on the *S. cerevisiae* pangenome, we selected the accessory ORFs that were previously detected in each of our 8 strains and selected the ones that had at least 1 read mapped on in both RNA-seq and Ribo-seq data in at least one of the replicates. We normalized the data as described previously using the inbuilt and default normalization in the *anota2seq* but by keeping genes with 0 reads values (Table S10, S11). The remaining ORFs were considered as expressed.

We first focused on the CPI isolate ORF acquired through introgression events (with the *Saccharomyces paradoxus* species). We selected the ORFs known to have an ortholog in *S. cerevisiae* genes. Then using homo/heterozygosity data (adapted from (20) gene presence/absence data), we selected the ORF that were homozygous for the *S. paradoxus* allele (n = 13) and we compared their expression with their orthologs in *S. cerevisiae*. We thus compared the introgression normalized values vs. their orthologs mean normalized value (obtained from the 7 other strains) using both RNA-seq or ribo-seq data.

We then explored the expression levels of the BPL isolate accessory ORFs, especially the ones acquired through horizontal gene transfer (HGT) by comparing their expressions (RNA-seq and Ribo-seq) and TE (Translation Efficiency, computed as the normalized Ribo-seq data divided by the normalized RNA-seq data) with the other BPL gene values. Similarly, we only focused on the HGT that were detected as expressed in at least one replicate. We computed a translation efficiency value by using the mean ratio of the ribo-seq normalized data divided by the RNA-seq normalized data.

## Data availability

All sequencing reads are available in the Gene Expression Omnibus (GEO) under the accession number GSE173654.

https://www.ncbi.nlm.nih.gov/geo/query/acc.cgi?acc=GSE173654

## Acknowledgments

We would like to thank Anne Lopes for helpful discussions on the manuscript. We are grateful to all the members of the Iwasaki laboratory for constructive discussions and technical help. This work was supported by a European Research Council (ERC) Consolidator grant (772505) to J.S. And E.T. was supported by the PhD Joint Programme CNRS & Weizmann Institute and a fellowship from the medical association la Fondation pour la Recherche Médicale (FDT202204014796). J.S. is a Fellow of the University of Strasbourg Institute for Advanced Study (USIAS) and a member of the Institut Universitaire de France. S.I. was supported by the Ministry of Education, Culture, Sports, Science and Technology (MEXT) (a Grant-in-Aid for Transformative Research Areas [B] “Parametric Translation”, JP20H05784), the Japan Society for the Promotion of Science (JSPS) (a Grant-in-Aid for Young Scientists [A], JP17H04998; a Challenging Research [Exploratory], JP19K22406), AMED (AMED-CREST, JP20gm1410001), RIKEN (Pioneering Project “Biology of Intracellular Environments” and Ageing Project), and the Takeda Science Foundation. Y.S. was supported by JSPS (a Grant-in-Aid for JSPS Fellows, JP19J00920) and RIKEN (Special Postdoctoral Researchers and Incentive Research Projects). Computations were supported by the supercomputer HOKUSAI Sailing Ship in RIKEN. Y.S. was a recipient of a JSPS Research Fellow (PD) and the RIKEN Special Postdoctoral Researchers Program.

## Notes

### Competing Interest Statement

The authors have declared no competing interest.

